# The application of drones for mosquito larval habitat identification in rural environments: a practical approach for malaria control?

**DOI:** 10.1101/2020.08.05.237933

**Authors:** Michelle C Stanton, Patrick Kalonde, Kennedy Zembere, Remy Hoek Spaans, Christopher M Jones

**Author notes:** Corresponding author: Michelle C Stanton, Vector Biology Department, Liverpool School of Tropical Medicine, Liverpool UK L3 5QA.

## Abstract

1. Spatial and temporal trends in mosquito-borne diseases are driven by the locations and seasonality of larval habitat. One method of disease control is to decrease the mosquito population by removing habitat and/or reduce the likelihood of larvae developing into adults, known as larval source management (LSM). In malaria control, LSM is currently considered impractical in rural areas due to perceived difficulties in identifying target areas. High resolution drone mapping is being considered as a practical solution to address this barrier. In this paper, we use our experiences of drone-led larval habitat identification in Malawi to assess the accuracy and practicalities of this approach.
2. Drone imagery and larval surveys were conducted in Kasungu district, Malawi between 2018-2020. Water bodies and aquatic vegetation were identified in the imagery using both manual methods and geographical object-based image analysis (GeoOBIA) and the performance of the classifications were compared. Larval sampling sites were characterised by biotic factors visible in drone imagery (e.g. vegetation coverage, type), and generalised linear mixed models were used to determine their association with larval presence.
3. Imagery covering an area of 8.9km^2^ across eight sites was captured. Characteristics associated with rural larval habitat were successfully identified using GeoOBIA (e.g. median accuracy = 0.98, median kappa = 0.96 using a standard RGB camera), with a median of 18.3% being classed as surface water, compared to 20.1% using manual identification. The GeoOBIA approach, however, required greater processing time and technical skills. Larval samples were captured from 326 sites, and a relationship was identified between larval presence and vegetation (log-OR=1.44, p=0.01). Vegetation type was also a significant factor when considering late stage anopheline larvae only.
4. Our study demonstrates the potential for drone-acquired imagery as a tool to support the identification of mosquito larval habitat in rural areas where malaria is endemic. There are, however, technical challenges to overcome before it can be smoothly integrated into malaria control activities. Further consultations between experts and stakeholders in the fields of drones, image analysis and vector control are needed to develop more detailed guidance on how this technology can be most effectively exploited.

## Introduction

Malaria cases in Africa have reduced by over half in the last two decades making transmission more heterogeneous. This has led to a growth of studies applying spatial and temporal analyses to determine where and when remaining transmission foci exist (Stresman, Bousema, & Cook, 2019), and a focus on how new and existing control methods can be best utilised to reduce this residual transmission (Bousema et al., 2016; Hsiang et al., 2020; Sy et al., 2019).

The geographical spread and extent of malaria transmission is limited by the seasonally-driven mosaic of water bodies available for female mosquitoes in which to lay their eggs. The ecology of preferred breeding grounds for mosquito oviposition vary both within and between species. For example, two of the main sibling species of the *Anopheles gambiae* sensu lato complex, *An. gambiae* and *An. arabiensis*, are found in transient, sunlit, small pools whereas *Anopheles funestus* is associated with more permanent, larger vegetated water (Nambunga et al., 2020). At the micro-geographic scale, the presence of mosquito larvae may differ over the course of just a few metres (Eneh, Fillinger, Borg Karlson, Kuttuva Rajarao, & Lindh, 2019; Gowelo et al., 2020; Musiime et al., 2020). Biotic and abiotic factors such as the presence of specific types of vegetation, microbiota, predators, algal density, shade, and water depth influence larval development.

Mosquito larval populations are fixed in space for the duration of their development to adulthood. Typically, eggs hatch into larvae within 2-3 days of oviposition and take 5-10 days to metamorphosise into pupae, although the speed of this process is highly dependent on temperature (Beck-Johnson et al., 2013). One method of controlling diseases transmitted by mosquitoes is to reduce the population by reducing the availability of oviposition sites and/or reduce the likelihood that resulting larvae develop into the adult stage (World Health Organization, 2013). Larval source management (LSM) involves the environmental, biological or chemical manipulation of the environment in which mosquitoes are present for the purpose of targeting the immature, aquatic stages of the mosquito and hence reducing the adult mosquito population. In the early days of mosquito control, an aggressive approach to searching and removing mosquito breeding sites was successful at reducing (and even eliminating) disease, with historical examples including its use during the construction of the Panama Canal in the early 20^th^ Century, and its role in the elimination of *Anopheles gambiae* in Brazil by 1940 (Tusting et al., 2013). In sub-Saharan Africa, LSM was responsible for large reductions in malaria incidence in Zambia copper mines between 1929 and 1949 (Fillinger & Lindsay, 2011). LSM is, however, a labour-intensive exercise and following the introduction of IRS by DDT in the 1950s and subsequently the development of ITNs in the 1990s, it fell out of favour as a viable control option, particularly in Africa where the long rainy seasons produce countless sites for *Anopheles* development (Fillinger & Lindsay, 2011). As such, LSM is currently only recommended as a complementary vector control intervention to bed nets and IRS to target residual transmission and as a method of combating insecticide resistance (Killeen, 2014). While its value is acknowledged by WHO and national malaria control programmes (NMCPs) there are several barriers to its widespread implementation.

The primary barrier to implementing LSM is the issue of determining where and when the intervention should be implemented. In rural settings the WHO recommend the application of LSM in areas where there is high coverage of long-lasting insecticidal nets (LLINs), evidence of outdoor biting and/or insecticide resistance and where larval sources are ‘*few, fixed and findable*’. Despite the lack of a clear definition of what can be considered ‘few’ or ‘findable’, this has led to many considering LSM to be impractical in rural areas with diffuse seasonal larval habitats. The perception of these terms may evolve as technology and processes for implementing LSM advance. This paper focuses on challenging the ‘*findable*’ component of this trio of conditions.

Geospatial technology is rapidly evolving and what now constitutes as ‘*findable’* may switch from less reliance on exhaustive ground-based searches to remotely sensed data. Drone mapping is being touted as at least equivalent (if not superior) to and more cost-effective than mapping larval habitat manually (Carrasco-Escobar et al., 2019; Hardy, Makame, Cross, Majambere, & Msellem, 2017) or using remotely sensed satellite imagery. While the latter can cover vast areas in a single day, images are often obscured by clouds and although very high-resolution commercial satellite imagery exists, the resolution (at best 30 cm) is still inferior to that obtained by drones (2-10 cm) with the time of image captured out of the data user’s control.

In this paper we explore the use of drones as a method for collecting very high resolution, contemporary imagery of an area for the purposes of identifying larval habitat. We tackle issues relating to the process of capturing drone imagery (by who, how much, how often), processing the images to extract the required information (what software, image classification methods, computer processing requirements), collecting ‘ground-truth’ data (entomological sampling), and subsequently summarising this information into recommendations that can be used by the control program implementers to guide their LSM activities.

## Materials and Methods

### Drone image capturing

A series of image data capture exercises were conducted within Kasungu district, central Malawi in an area that has been designated by the Government of Malawi, in collaboration with UNICEF, as a ‘humanitarian drone testing corridor’ (Fig. 1). Authorisation to conduct these flights was obtained from the Malawi Department of Civil Aviation. Malaria transmission occurs all year round in this area, with parasite prevalence in children between 2 and 10 years old estimated at 19% in 2017 (Chipeta et al., 2019). This transmission is potentially driven by a number of reservoirs which provide permanent sources of water within which female *Anopheles* can lay their eggs (Kibret, Lautze, McCartney, Nhamo, & Wilson, 2016). Images were captured over three visits in June 2018 (early dry season), October 2019 (late dry season) and February 2020 (wet season) using two drones, both of which were able to capture images using a standard RGB camera, plus a near-infrared (NIR) camera. Both drones were purchased off-the-shelf from commercial vendors. The first was a multirotor (quadcopter) type aimed towards the ‘hobbyist’ market (the DJI Phantom 4 Pro), supplemented by an additional NIR sensor by Sentera (Sentera, 2020). The second was a fixed-wing drone marketed towards the agriculture industry (eBee SQ) which incorporated a Parrot Sequoia multispectral camera.

**Figure 1:**
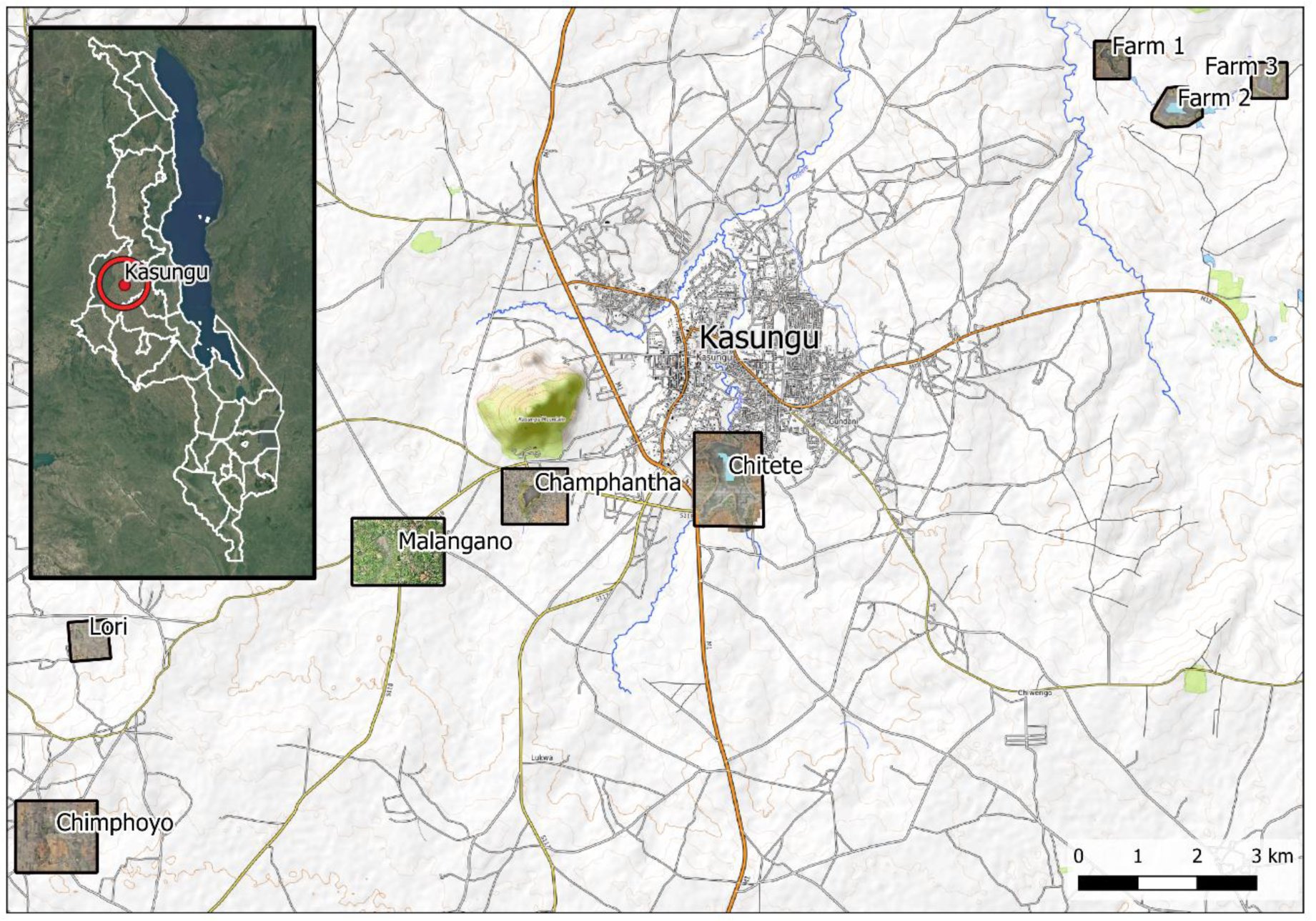
Locations of sites surveyed within the ‘humanitarian drone testing corridor’, centred on Kasungu town, Central Malawi (inset). Coordinates can be found in Table S1.

### Drone image processing

Individual images captured during each mapping mission were stitched together into orthomosaics using the commercial image processing software Agisoft Metashape Professional (version 1.4.2). A subset of images captured during the wet season were classified using a geographical object-based image analysis (GeoOBIA), using the LargeScaleMeanShift algorithm within the open source software Orfeo Toolbox (version 7.1.0), applied within the QGIS environment (version 3.8.1). GeoOBIA involves grouping contiguous pixels into ‘objects’ or ‘segments’ such that each segment is relatively homogenous (within a prespecified threshold) with respect to pixel characteristics. In this instance, pixels were grouped into segments according the values of red, green, blue and elevation, with the latter being estimated using photogrammetric methods within Agisoft Metashape and then rescaled to lie between 0-255 to match the scale of the RGB values. We used trial and error to select the optimal segmentation parameters i.e. the spatial radius, range radius and minimum segment size. The smoothing radius determines how the amount by which the image is smoothed or filtered prior to the segmentation algorithm being implemented, whereas the range radius determines the similarity between pixels for grouping within the same segment. Similarity in this context refers to the Euclidean distance between two pixels. Supervised classification was then undertaken to assign each segment to one of 12 land cover classes (Table S2), including open water and aquatic vegetation (floating, emerging, submerged) based on the characteristics of that segment. Table S3 displays the segment-level characteristics used, which incorporated characteristics related to segment texture (Haralick textural features (Haralick, Dinstein, & Shanmugam, 1973)) in addition to a range of water and vegetation indices. The mean and variance of each of these were used in the classification.

Classification was undertaken using a set of 1800 segments all of which were firstly manually classified by the research team. One-third of the segments (n = 600) were within a 400m by 400m area and were used to train the classification algorithm. An additional one-third were in the same 400m by 400m area and were used for evaluating the accuracy of the classification within the same geographical area used for training (spatial interpolation), whereas the remaining 600 segments were distributed outside of the area used for training (spatial extrapolation). Classification was undertaken in R (version 3.6.1) using the caret package and the Random Forests classification algorithm (Kuhn, 2008). Manually classified segments within the 400m by 400m area were randomly split into training and testing segments. A ten-fold cross validation approach was used to determine the optimal tuning parameters for the Random Forests algorithm, and the resulting model was then applied to the testing segments. An accuracy statistic (% of classifications that were correct), and the Cohen’s kappa agreement statistic (Cohen, 1960) were then calculated for the interpolated and extrapolated testing segments initially considering all 12 land cover classes, followed by a reduced classification that only differentiated between surface water (open water and aquatic vegetation) and any other class. The classification was then applied to the entire 600m by 800m area, and the percentage of the area classed as being covered in surface water was calculated. This process of randomly splitting the segments into training and testing groups was repeated 100 times, and the median and inter-quartile range of the resulting accuracy, kappa agreement and percentage surface water cover were reported.

A manual classification of surface water was also undertaken which involved systematically scanning through the image and creating a polygon around each area of surface water. The surface area of the image manually identified as being covered in water was then compared with that classified as surface water using the automated approach. The time taken and the computer resources required for each of these tasks were also recorded.

An overview of the image capture, processing and classification procedures is presented in Fig. 2.

**Figure 2:**
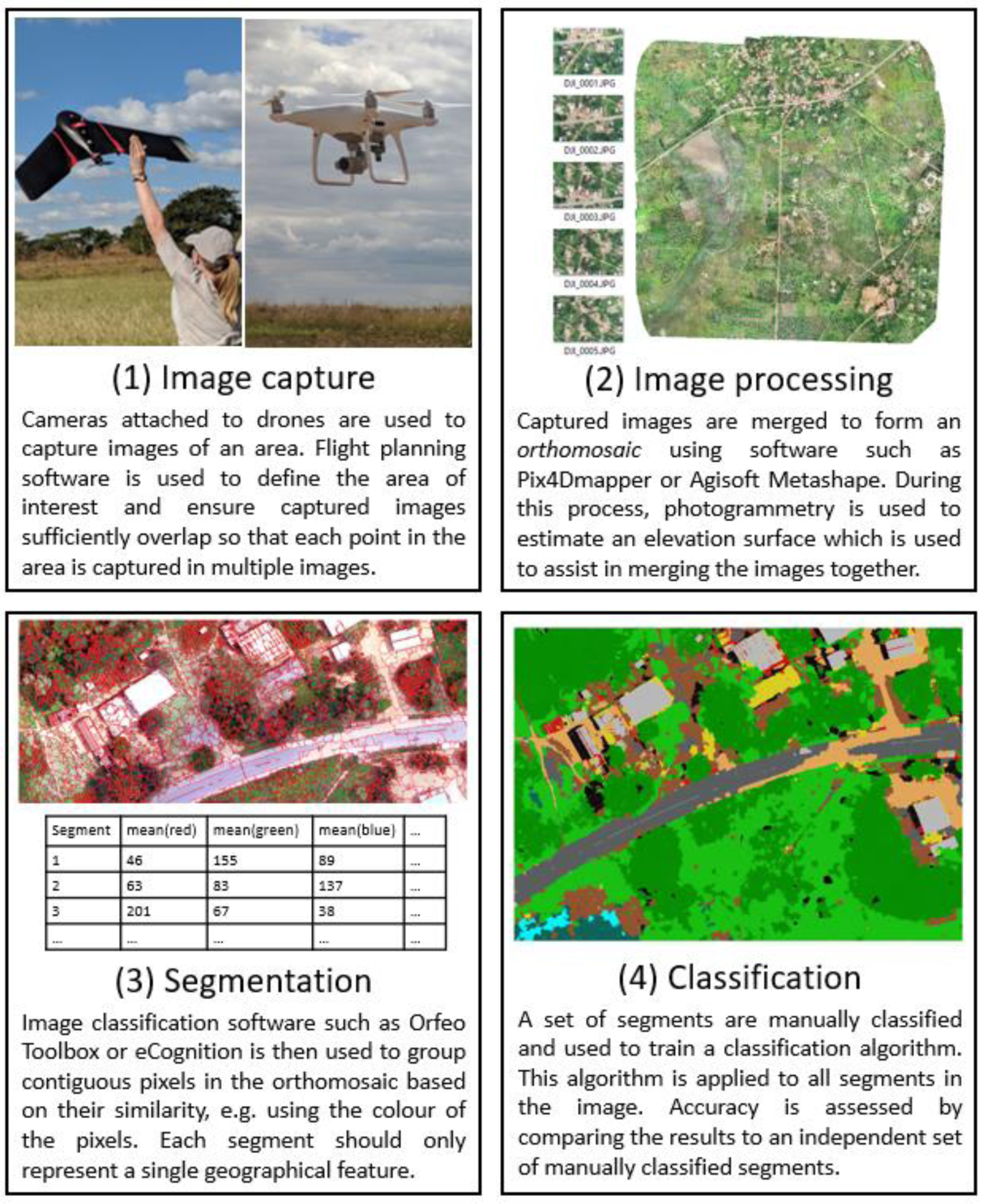
Processes undertaken to identify larval habitat from drone imagery.

### Entomological sampling

Larval surveys were conducted concurrently to the drone image capture during each field visit (Fig. 1). Permission to collect these data were obtained from the Kasungu District Council, Kasungu Water Board and private landowners. As the first two field visits were undertaken during the dry season, sampling was focused in and around permanent water bodies which in this case were local reservoirs that provide drinking water and irrigation to the area. Sampling was undertaken at regular intervals around the periphery of reservoirs. Reservoirs were selected purposively based on their proximity to Kasungu town (for accessibility) and their proximity to human settlements. The third visit was conducted during the wet season, and sampling was focused around one of the reservoirs sampled in the previous dry season. Both temporary and permanent water bodies were sampled, with sampling sites identified from drone imagery captured the previous day. A subset of sites was sampled on four consecutive days to determine their consistency with respect to larval presence.

At each site, the presence and number of larvae were recorded using 10 repeated dips of the surface water, categorised by stage (L1/L2 or L3/L4) and either anopheline or culicine. To characterise malaria vectors in Kasungu as part of our broader efforts to understand transmission in the area, we raised all anopheline larvae to adult stage and identified morphologically to species (Coetzee, 2020). The location, description and photographs of each site were recorded using an Android Smartphone and Open Data Kit (ODK). These photographs were later used to classify each site according to the amount and type of vegetation present plus turbidity. A generalised linear mixed model was then fitted to the resulting presence/absence data to predict the likelihood that larvae were present from biotic site information obtained via drone imagery (vegetation type, coverage, turbidity). Models were fitted to presence/absence data for any larvae irrespective of stage or genus, and for late stage larvae only as characteristics of habitat containing late stage larvae are considered by WHO to be of greater importance than early stage (World Health Organization, 2013). The productivity of the sampled sites with respect to the number of early or late stage larvae collected was also considered.

## Results

### Image capturing

During the three sampling periods we captured a total of 10 distinct areas in Kasungu, covering an area of 8.9km^2^. The two drones significantly differ in relation to operational costs, equipment and software requirements and usage. Tables 1 and 2 describe the primary differences in relation to initial costs and operational usage respectively. The fixed wing drone (eBeeSQ) had a greater initial cost than the multirotor Phantom 4 Pro due to it being inclusive of a NIR sensor, costing approximately £7000 (inclusive of an educational discount), compared to £3,300 for a standard (RGB sensor) Phantom 4 Pro drone on which a NIR sensor was retrofitted. The eBeeSQ also required a high-spec laptop (£1000+) on which to run the software required to plan and conduct missions, whereas the Phantom 4 Pro was operated using free apps installed on GPS-enabled Android or iOS smartphone or tablet devices. On an operational level, the primary differences are between the flight times per battery, and the ease of use (Table 2). While, overall the Phantom 4 Pro is easier to use due to the small amount of open space required for take-off and landing, its limited battery life means that to cover a relatively modest area of 1km^2^, 2-3 individual flights are needed depending on whether a fixed launch site is used or whether this is adapted to minimise flight time. This comes at a cost of both time and money, particularly given each battery comes at a price of £150. The eBeeSQ fixed wing drone requires less energy to fly and therefore batteries last approximately twice as long (up to 1 hour in comparison to 30 minutes) than the Phantom 4 Pro. Therefore, while the time required to cover 1km^2^ is longer (66 minutes compared to 38 minutes when flown at 120m above sea level [asl] which an 80% overlap in captured images), this area can be comfortably covered using two batteries and a single launch site, meaning that in practice the process is more efficient. The fixed-wing drone is however more difficult to operate than the Phantom 4 Pro, requires a larger space for take-off and landing, and therefore cannot be used in more densely vegetated areas.

**Table 1:** Comparison of approximate costs required to capture and process imagery captured by the Phantom 4 Pro (rotor) and eBeeSQ (fixed wing).

|  |  |  | **Phantom 4 Pro** | **GBP (£)** | **eBeeSQ** | **GBP (£)** |
| --- | --- | --- | --- | --- | --- | --- |
| Costs | Initial costs | Standard drone | RGB only | £1500 | RGB only | N/A |
|  |  | Supplementary sensor | Sentera NDVI | £1800 | Parrot Sequoia | £7000+ |
|  | Supplementary hardware |  | Tablet | £150 | High spec laptop | £1000+ |
|  |  | Spare batteries | <=30 mins flight time (per battery) | £150 | <=1hr flight time (per battery) | £90 |
|  | Supplementary software | Mission planning | Pix4D capture | £0 | eMotion Ag | £0 |
|  |  | Image processing | Agisoft MetaShape Professional Edition (Educational Licence) | £425 | Agisoft MetaShape Professional Edition (Educational Licence) | £425 |
|  |  | Image classification | Orfeo Toolbox | £0 | Orfeo Toolbox | £0 |

**Table 2:**
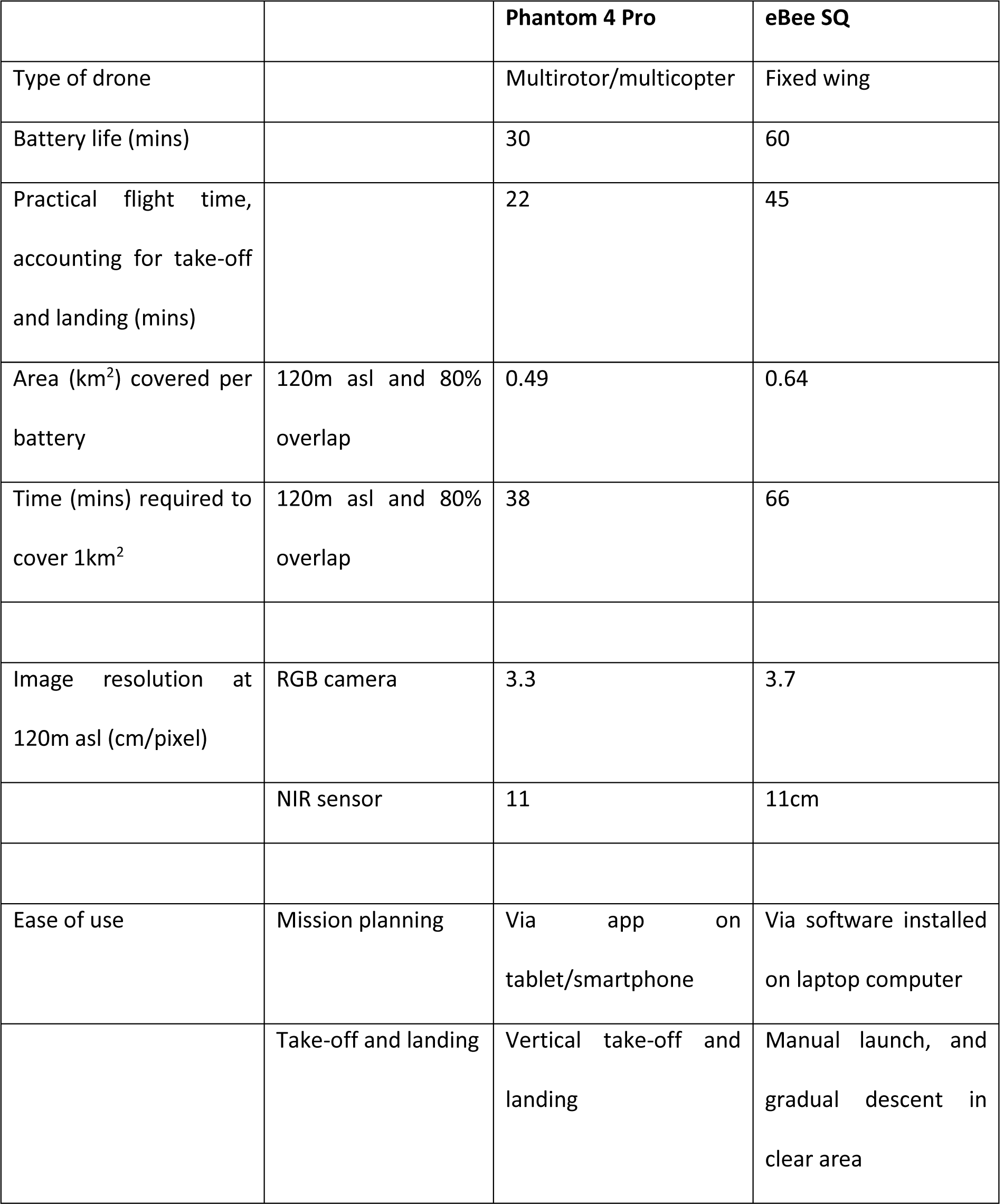
Practical and operational differences between the drones used in this study

### Image processing & classification

We used Agisoft Metashape to process all images captured. Using the Phantom 4 Pro flying at 120m above surface level, with an 70% overlap in images, a total of 782 individual images (6.2GB) covered an area of 1.77km^2^. Processing these images in Agisoft Metashape in order to produce an orthomosaic of the area and an accompanying digital surface model took a total of 250 minutes using a computer with an Intel Core i7-6700 processor, 32GB RAM, resulting in an orthomosaic with a spatial resolution of 3cm (file size = 4.2GB). A subset of the image covering an area of 0.48km^2^ (800m by 600m, file size = 1.9GB) was then selected for classification (Fig. 3).

**Figure 3:**
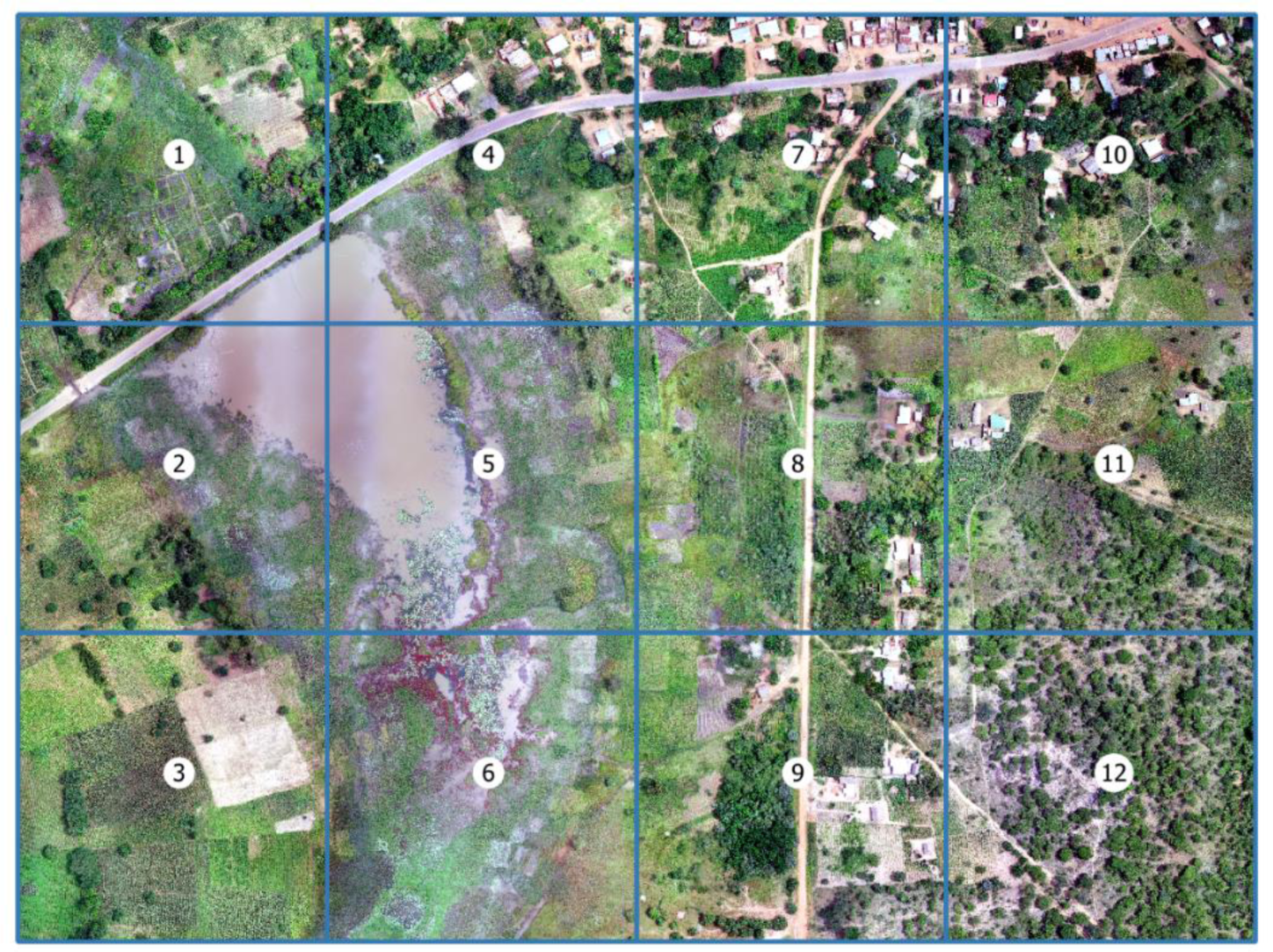
Image captured by the Phantom 4 Pro in Kasungu in February 2020 covering 800m by 600m, with each grid representing 200m by 200m. Grids 4, 5, 7 and 8 were used for training the classification algorithm, and an assessment of its accuracy was made using features both within this area (interpolation) and in the surrounding grids (extrapolation).

A set of 1800 training and testing segments were then generated for the 12 identified land classes (Table S2), plus an additional category representing areas that were in shadow. The study area was partitioned into cells of 200m by 200m, labelled as cells 1-8 (Figure 3), and the training and testing segments were proportionally distributed throughout the cells as follows: two thirds (1200) of segments were within cells 4, 5, 7 & 8 covering an area of 400m by 400m. We refer to these as the *internal* segments. One third of segments (600) were within the remaining cells (1-3, 6, 9, 10-12) which we refer to as *external* segments.

Segmentation was performed using Orfeo Toolbox (OTB) functions within the QGIS environment. Figure 4 demonstrates the impact of varying values of the spatial and range radius on the resulting segmentation and the time taken to perform this segmentation over a 100m by 100m area using the computer specifications previously specified (see Methods). While increasing the spatial radius provided a more adequate balance between over-segmentation (single discrete features of interest being split into many segments) and under-segmentation (multiple discrete features of interest being grouped into a single segment), this came at the price of substantially increasing the processing time. Additional processing time is required to calculate the segment-level summaries (mean, variance) of each of the variables being used to classify the imagery, with processing time increasing as the number of segments increases.

**Figure 4:**
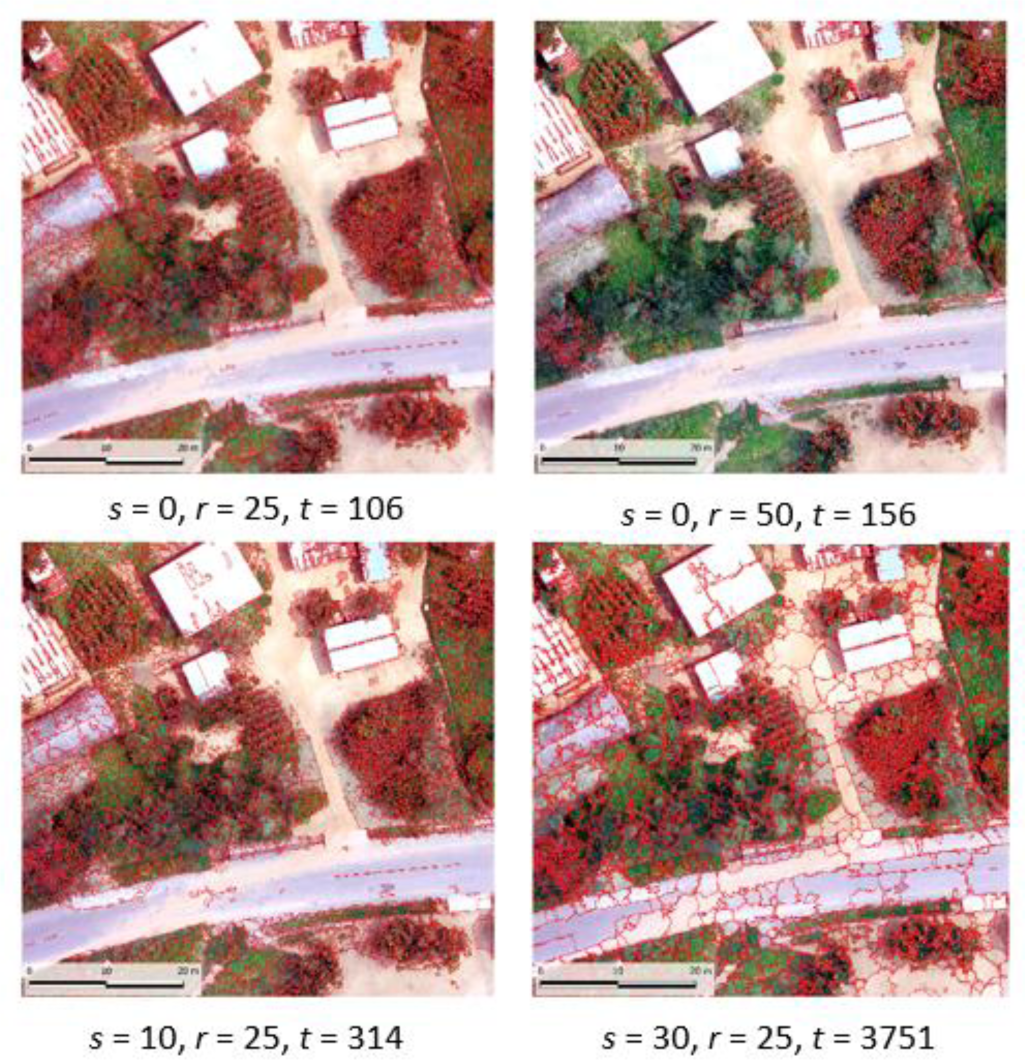
Examples of the segmentation process under different values for spatial radius *s* (0, 10, 30), range radius *r* (25, 50) with a minimum segment size of 100. Time *t* corresponds to the time taken in seconds to segment a 100m by 100m image with a spatial resolution of 3cm using the LargeScaleMeanShift algorithm in Orfeo Toolbox. This process includes calculating the mean and variance of the RGB and elevation values for each segment.

The segmentation process was then applied to the entire 800m by 600m area using the parameters 10 (spatial radius), 25 (range radius) and 200 (minimum segment size), creating close to 800,000 segments. The total processing time, which includes calculating the segment-level mean and variance of the RGB and elevation values, was 24.5 hours with an additional 15 hours taken to calculate the mean and variance of each of the additional variables under consideration (Table S2). Two classifications were then undertaken, one of which included the NIR-derived variables and one of which did not.

The resulting accuracies of these classifications are presented in Table 3, and a representation of the classified output from one area of Kasungu excluding NIR-derived variables is shown in Fig. 5, with the resulting image which includes NIR-derived variable presented in Figure S1. The corresponding variable importance plots are available in the SI (Figure S2). The results are very similar for both models fitted with and without the NIR-derived variables. The variable making the greatest contribution to the classification model in both cases is the mean elevation, with mean red, blue, green and brightness also important. While the NIR-derived variables NDVI and SAVI make the greatest contribution to the classification algorithm in the second model (Figure S2), Table 3 indicates that the inclusion of NIR-derived variables does not make any significant impact on classification accuracy, with overall median interpolated accuracy obtained using NIR-derived variables being marginally lower (0.904) than that obtained without using NIR-derived variables (0.910). There is a clear drop in both accuracy and kappa agreement when considering data from the extrapolation area, with accuracy reducing to 0.761 and 0.798 when considering overall accuracy without and with NIR-derived variables respectively. This reduction in classification quality is less pronounced when considering surface water (open water or aquatic vegetation) accuracy alone compared with trying to distinguish between all 12 land cover classes. Trends in values of kappa are similar.

**Table 3:** Summaries of classification accuracy (proportion of segments correctly classified) and kappa agreement for all 12 classes (*overall)* and for *surface water* (including open water and aquatic vegetation) versus all other classes for GeoOBIA obtained with and without NIR-derived variables.

|  | Area | Without NIR-derived variables |  | With NIR-derived variables |  |
| --- | --- | --- | --- | --- | --- |
|  |  | Median | IQR | Median | IQR |
| Overall accuracy | Interpolated | 0.910 | 0.898-0.917 | 0.904 | 0.899-0.908 |
|  | Extrapolated | 0.761 | 0.754-0.772 | 0.798 | 0.758-0.811 |
| Surface water accuracy | Interpolated | 0.983 | 0.980- 0.986 | 0.982 | 0.979-0.986 |
|  | Extrapolated | 0.942 | 0.941-0.947 | 0.936 | 0.931-0.939 |
| Overall kappa | Interpolated | 0.902 | 0.888-0.909 | 0.895 | 0.890-0.900 |
|  | Extrapolated | 0.738 | 0.731-0.750 | 0.779 | 0.735-0.793 |
| Surface water kappa | Interpolated | 0.960 | 0.954-0.967 | 0.958 | 0.950-0.967 |
|  | Extrapolated | 0.873 | 0.868-0.881 | 0.855 | 0.844-0.864 |
| % surface water | All | 18.3 | 17.3-20.9 | 16.6 | 15.7-22.0 |

**Table 4:**
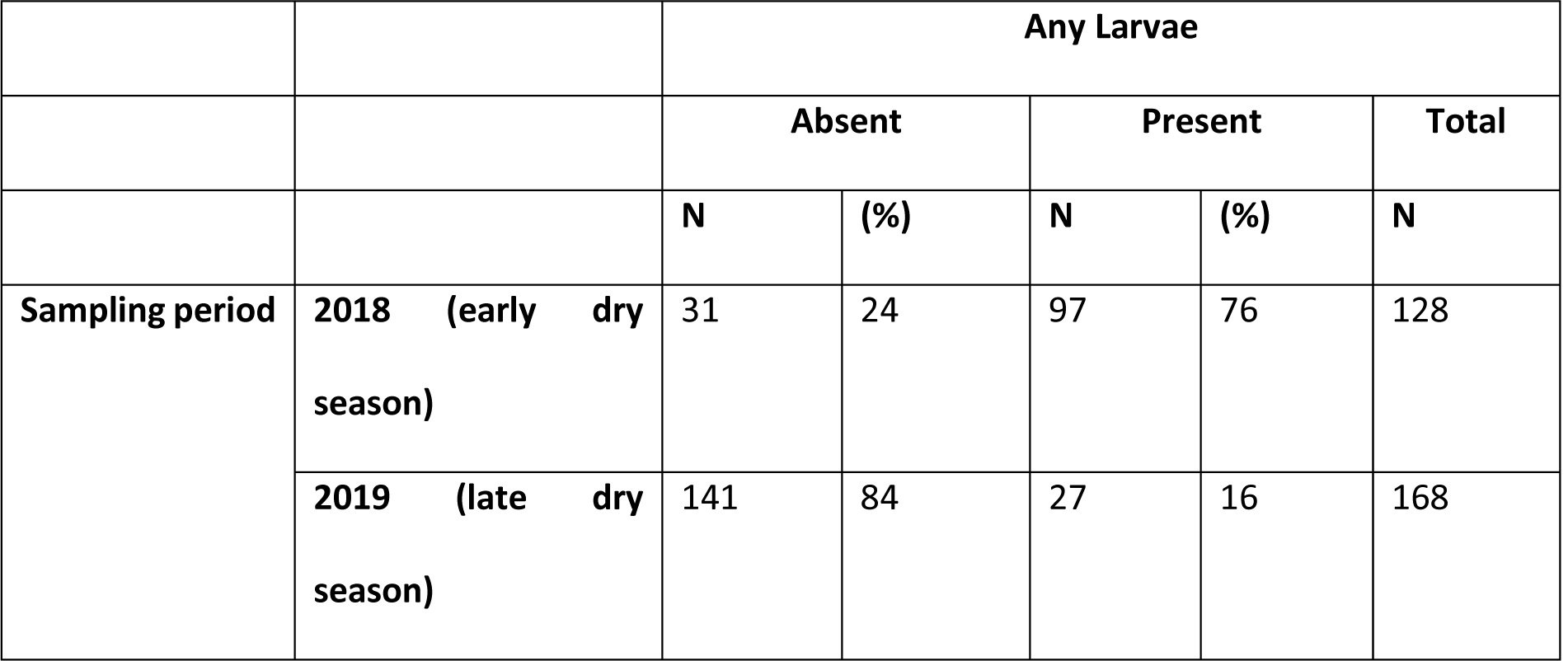
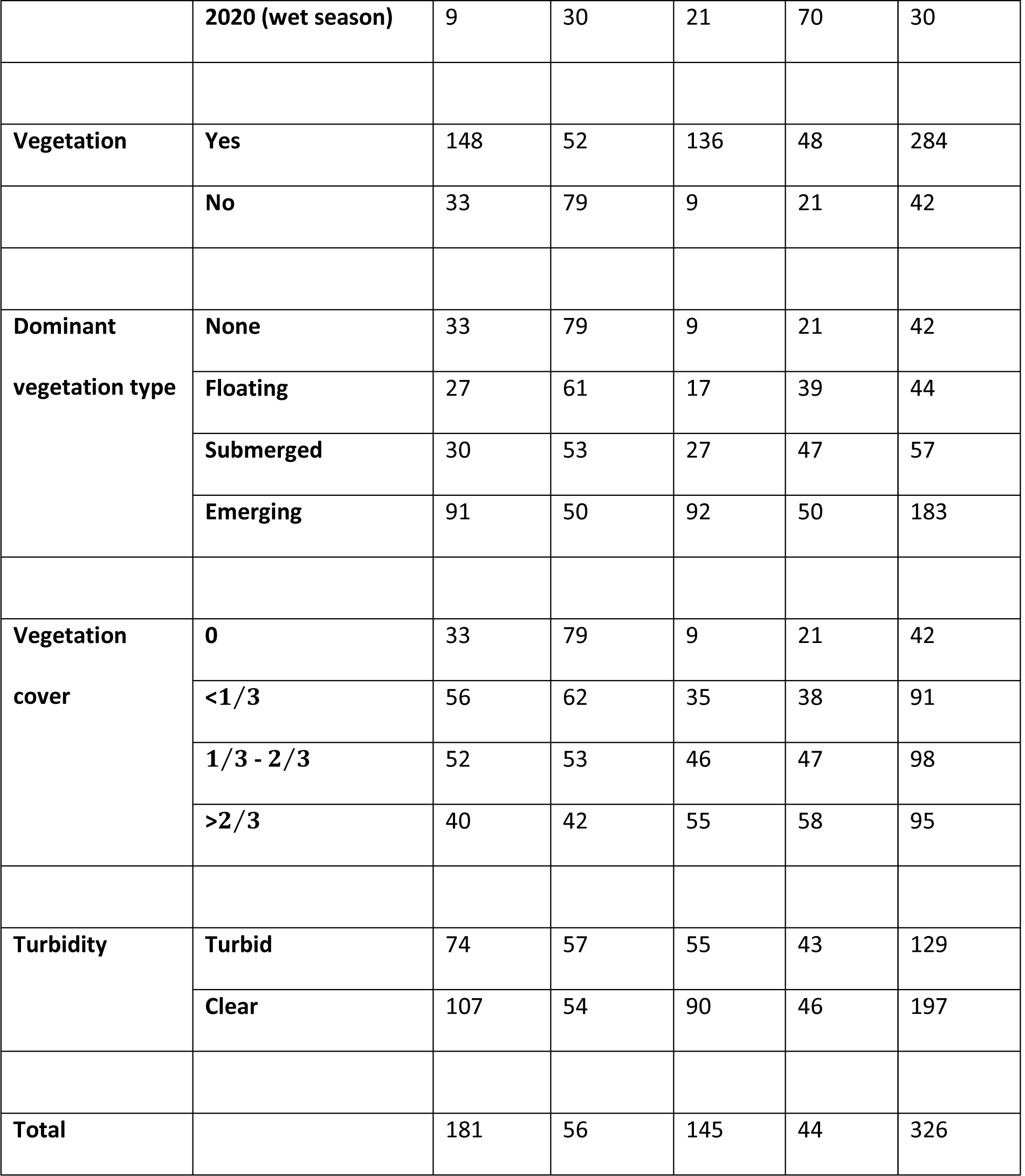
Summaries of the larval sampling sites by presence/absence of mosquito larvae found. A similar table for late stage (L3-L4) larvae can be found in the SI (Table S4).

|  |  | Any Larvae |  |  |  |  |
| --- | --- | --- | --- | --- | --- | --- |
|  |  | Absent |  | Present |  | Total |
|  |  | N | (%) | N | (%) | N |
| Sampling period | 2018 (early dry season) | 31 | 24 | 97 | 76 | 128 |
|  | 2019 (late dry season) | 141 | 84 | 27 | 16 | 168 |
|  | **2020 (wet season)** | 9 | 30 | 21 | 70 | 30 |
| **Vegetation** | **Yes** | 148 | 52 | 136 | 48 | 284 |
|  | **No** | 33 | 79 | 9 | 21 | 42 |
| **Dominant**<br>**vegetation type** | **None** | 33 | 79 | 9 | 21 | 42 |
|  | **Floating** | 27 | 61 | 17 | 39 | 44 |
|  | **Submerged** | 30 | 53 | 27 | 47 | 57 |
|  | **Emerging** | 91 | 50 | 92 | 50 | 183 |
| **Vegetation**<br>**cover** | **0** | 33 | 79 | 9 | 21 | 42 |
|  | **<1/3** | 56 | 62 | 35 | 38 | 91 |
|  | **1/3 - 2/3** | 52 | 53 | 46 | 47 | 98 |
|  | **>2/3** | 40 | 42 | 55 | 58 | 95 |
| **Turbidity** | **Turbid** | 74 | 57 | 55 | 43 | 129 |
|  | **Clear** | 107 | 54 | 90 | 46 | 197 |
| **Total** |  | 181 | 56 | 145 | 44 | 326 |

**Figure 5:**
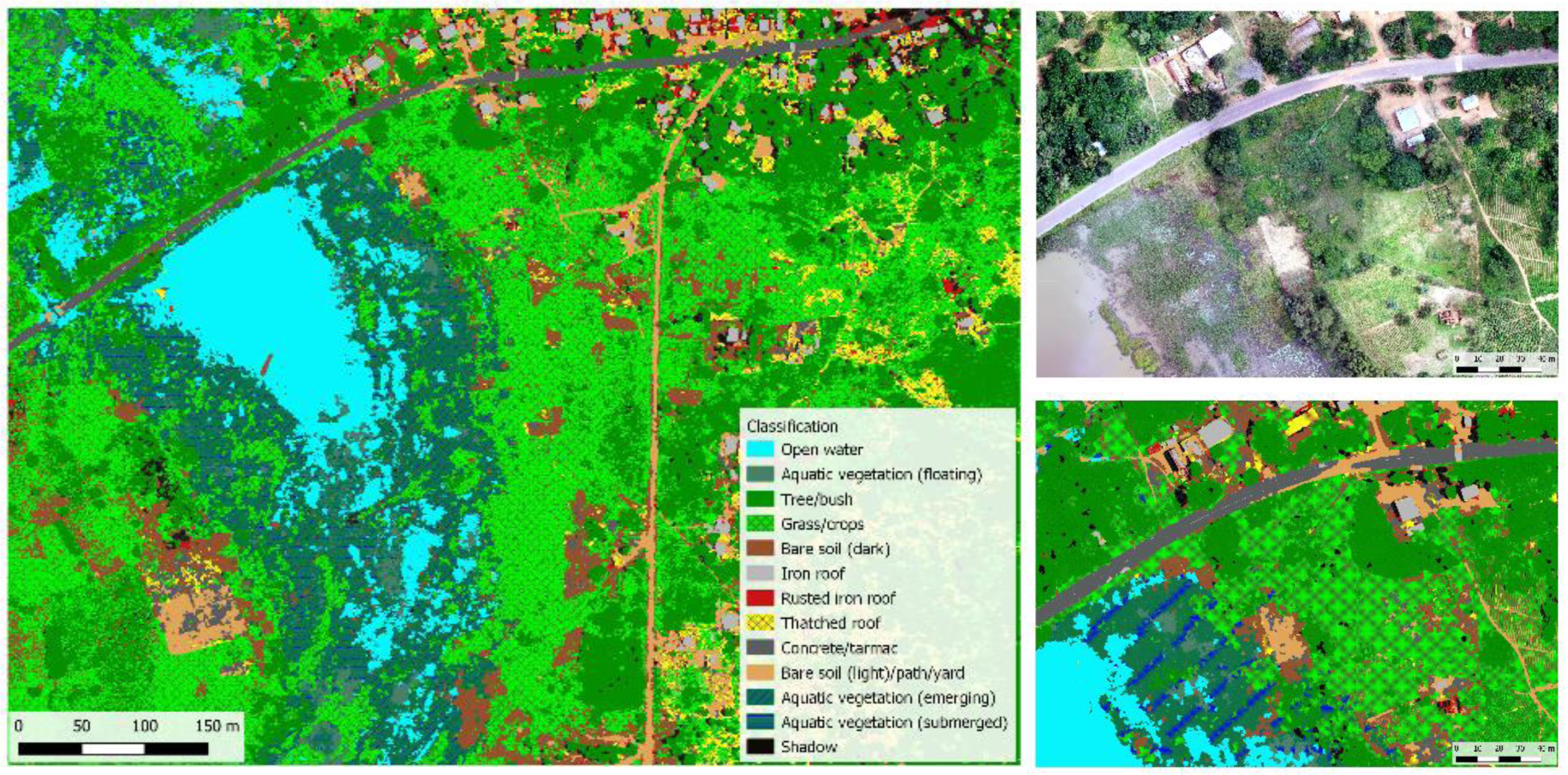
Example of a classification obtained for the entire study area using the random forests algorithm without including NIR-derived variables (left), with a more detailed view of a smaller area comparing the original image (top right) with the classified image (bottom right).

There is a close agreement in the percentage of the 400m by 600m area that is covered in surface water obtained by fitting the model with and without NIR-derived variables (without NIR: median = 18.3%; with NIR: median = 16.6%), with more variability observed in the NIR-inclusive models (Table 3). A manual review of the image, independently undertaken by two researches, resulted in a larger percentage of the area being identified as surface water (21.2% and 20.1% by researchers 1 and 2 respectively), with the intersection of these two outputs (our ‘*manual surface water classification’*) covering 20.1% of the area. This process of manual classification took approximately two hours to complete.

### Entomological surveys

During three separate field visits (June 2018, October 2019, and February 2020) a total of 326 larval sites were sampled (available through the Figshare repository (Stanton, 2020)). During the dry periods these samples were focused along the shorelines of larger permanent water bodies (296 sites), with a mixture of 30 temporary and permanent sites surveyed during the wet season. Both anopheline and culicine larvae were found throughout the area during each sampling period (56% of sites sampled), with the lowest proportion of positive sites found in the late dry season (76% in June 2018, 16% in October 2019, 70% in February 2020). No clear sympatry was observed between anopheline and culicine larvae in this study. For example, of the 321 sites where late stage larvae data were recorded (excluding 5 sites with missing data), larvae were observed in 31% (101) of samples with only 7% (24) sites containing both anophelines and culicines. In the area surrounding the Malangano site (Fig. 3) in February 2020, we morphologically identified 177 out of 297 anopheline specimens to species level, finding a predominance of *An. gambiae s*.*l*. (87.6%) followed by *An. coustani* (8.5%) and very few *An. pretoriensis* (2.3%) and *An. funestus* (1.7%).

At each site, GPS coordinates were recorded using the ODK app, photographs were taken using a smartphone, and aerial imagery was captured (Fig. 6). Samples were taken within approximately one metre of where the researcher stood to record the coordinate, however, as GPS coordinates have an accuracy of approximately three metres it was not possible to pinpoint precisely where the samples were taken within the aerial imagery.

**Figure 6:**
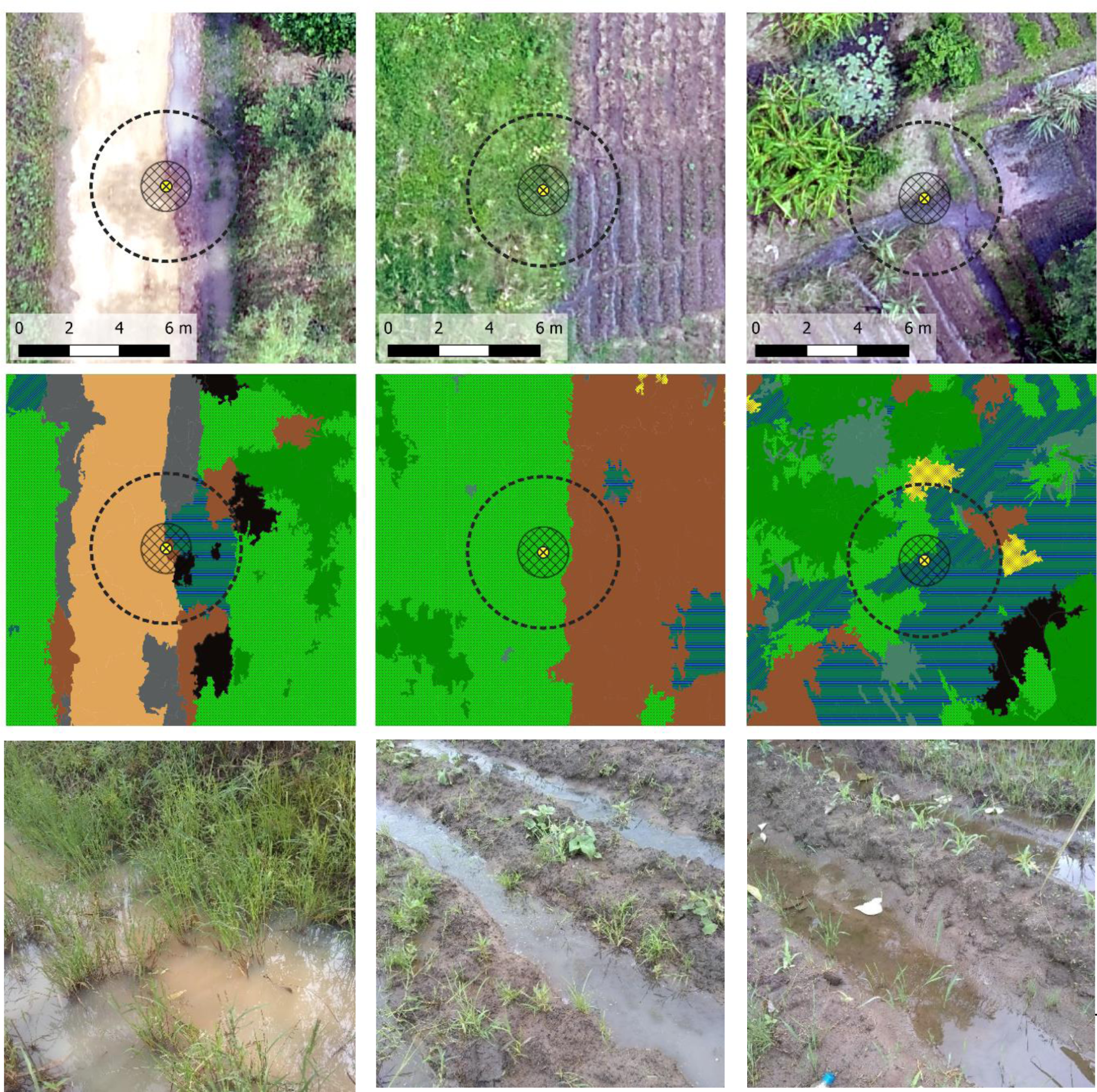
Examples of sampling sites where anopheline larvae were found. The top row indicates the precise GPS location captured using ODK (yellow circle), the expected sampling area based on these coordinates (1m radius), and the expected accuracy of the coordinates (3m radius), overlaid on top of the drone imagery. The middle row presents the classified imagery for these sites and the bottom row contains photographs of each site taken at the time of sampling.

Using the photographs, we characterised each site according to presence/absence of larvae and sample site characteristics including dominant vegetation type (none, floating, submerged, emerging), vegetation cover (none, <1/3, 1/3 - 2/3, >2/3) and turbidity. Vegetation was present in most sites sampled (284/326, 87%). Of these, 64% (183) contained emergent vegetation, 20% (57) contained submerged and 15% (44) contained floating vegetation. Vegetation cover varied evenly across sites, with 32% having low (0-1/3) coverage, 35% having moderate (1/3-2/3 coverage) and 33% having high (>2/3) coverage. There was an interaction between vegetation type and coverage, such that sites with floating vegetation rarely had high vegetation coverage. With regards to turbidity, 40% (129) of sites were classed as turbid whereas the remaining 60% (197) were clear.

A strong interaction was observed between vegetation type and coverage, therefore when fitting the GLMMs to the presence/absence data we did not consider these variables in the model simultaneously, but rather explored which of the two resulted in the best fitting model with regards to AIC. After counting for the effect of sampling period and site, there was a strong association between the presence of vegetation and the likelihood of any larvae (log-OR=1.44, p=0.01), however accounting for vegetation coverage or vegetation type did not improve the model further. When considering *Anopheles* L3 and L4 larvae only, the model was improved when vegetation type was considered such that larvae were more likely to be present when emerging (log-OR=1.14, p=0.07) or submerged (log-OR=1.90, p=0.01) vegetation were available, compared to sites with no vegetation. With regards to productivity, while there was variability in the abundance of larvae sampled per site (145 sites, min = 1, median = 4, max = 56), there were insufficient high productivity sites to formally explore any trends in their characteristics.

During the wet season, 10 sites were repeatedly sampled over four days, with larvae consistently observed in four sites and no larvae being found on at least one day in the remaining six sites. We observed that due to changes in the environment it was difficult to resample the same locations across larger time scales. Temporary surface water observed in the wet season dried up even after just a few days without rain and shorelines of permanent water bodies varied substantially both between seasons and between the same season over consecutive years (Fig. 7). For example, we observed that images captured later in the dry season (October) in 2019 were wetter than those captured in the early dry season (June) in 2018 at the Chitete reservoir.

**Figure 7:**
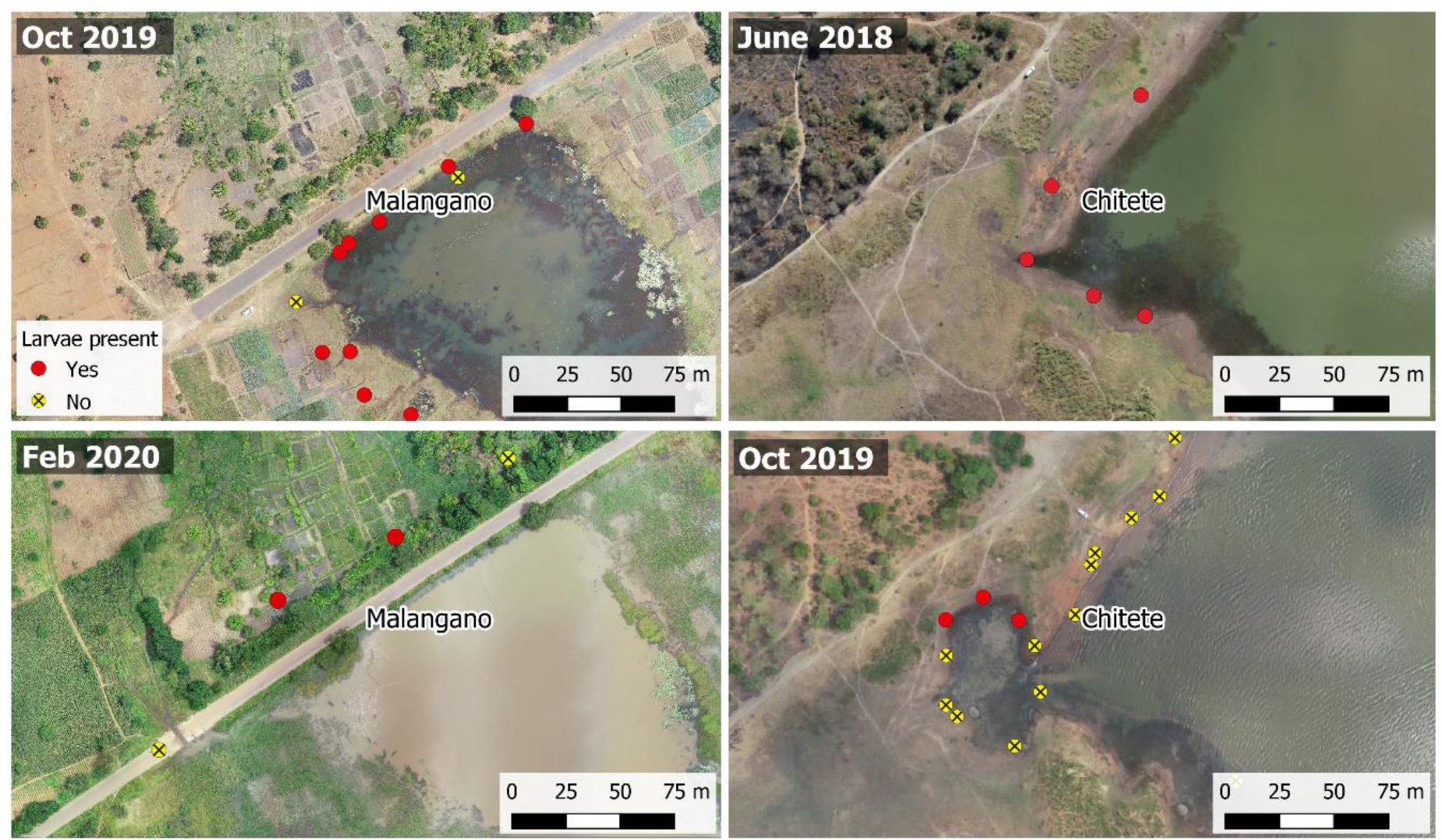
Comparisons of aerial images captured at different seasonal time points. Left images display a comparison between consecutive dry (Oct 2019) and wet (Feb 2020) seasons around the Malangano dam. Right images display comparisons between dry seasons over two consecutive years (June 2018, Oct 2019) around the Chitete dam.

## Discussion

### Image capturing

Image capture using drones inevitably leads to technical and skills-based challenges and we highlight a few of these here in the context of searching for water bodies in a rural setting. Aside from hardware and software issues we noted that flight experience was a key requirement to determine optimal flight times as neither the rotor or fixed wing drone could be flown in wet or windy conditions, and we experience the impact of extreme weather on the hardware with multiple occasions of over-heating on warm days. This highlighted the need for extensive drone piloting training by the operator. The country’s drone regulations also need to be taken into careful consideration. In Malawi, data capture was facilitated by the relationship between UNICEF and the Department of Civil Aviation and a toolkit is currently being developed to outline the procedures that need to be followed by those wishing to fly drones for non-commercial purposes (https://www.updwg.org/wp-content/uploads/2019/12/Malawi-RPA-Toolkit-2019_Dec.-Final.pdf). While regulations vary by country, national civil aviation authorities are also requiring drone pilots to obtain accredited qualifications and seek appropriate permissions before using drones for research or humanitarian purposes. Training courses which cover both the operational and regulatory aspects of drone flying are currently quite sparse in sub-Saharan Africa and this may require future pilots to travel outside of their own country to gain the necessary experience. Should a malaria control implementer wish to use drone imagery within their programmes, they may therefore incur significant expense both in purchasing the equipment and training their staff. A solution to this would be to outsource the image capturing to qualified drone pilots operating in the area.

An additional bottleneck is the availability of hardware within the country of operation. While it may be possible to purchase off-the-shelf drones in-country, should any technical issues arise, obtaining part replacements or repairs becomes problematic and expensive. Investments are therefore being made in ‘home-grown’ drones, to support local economies, decrease the cost of equipment, and make repairs much more easily accessible. In Malawi for example, MicroMek (https://www.micromek.net/) manufacture the low-cost fixed wing drone known as EcoSoar (Standridge, 2018), for both transporting goods and capturing imagery.

### Image processing

Processing drone imagery to create the orthomosaics is time-consuming, requires a high-spec computer and a large capacity for data storage. Therefore, to use this imagery in the field, an NMCP would require people skilled in both image capture and processing, plus access to the relevant software. These skills are not usually taught as part of standard drone pilot training, however this may change as the potential for using drone technology for humanitarian purposes is increasingly realised. For example, the African Drone and Data Academy was launched in January 2020 in Malawi to build capacity in both drone piloting and drone image processing and analysis (UNICEF, 2020).

Image classification is appealing because once the algorithm has been trained, it can simply be applied to any additional imagery captured without any or only a little additional data being required. In our analysis we showed that there was a decline in classification accuracy in areas within very close proximity to that used to train the algorithm and noted that even in this small area there were important land cover classes in the extrapolation area that did not appear in the training area e.g. red algae in the water. This challenge is likely to be exacerbated when considering areas further apart, or data collected at different time points. As more data are collected these limitations may be overcome, however in the short term, the effort required by the end-user e.g. an individual NMCP to train a classification algorithm may outweigh its benefits. In this demonstration we implemented a geographical object-based classification approach which generated the segments and computing segment-level characteristics prior to training and applying the classification algorithm. The segment-generating process can be very time-consuming depending on the values of the segmentation parameter referred to as the spatial radius, and the size of the area being classified. While other classification techniques such as pixel-based classification may be quicker to perform, an object-based approach is the most appropriate for very high-resolution images such as that generated by drones (Pande-Chhetri, Abd-Elrahman, Liu, Morton, & Wilhelm, 2017). The cost and benefits of accuracy against processing time therefore need to be considered should an NMCP wish to perform image classification in-house. The role of additional sensors in the image classification process is also unclear. In this analysis we compared the classification accuracy using imagery captured from a standard camera only (RGB), plus additional imagery captured by a much more expensive NIR sensor. While our performance metrics indicated very little difference in the accuracy obtained using the two approaches, a more extensive investigation would need to be undertaken over a more environmentally diverse area before we can conclude whether or not there is a benefit to incorporating this additional technology.

A more practical solution to ‘automated’ image classification may be to persevere with the less efficient, but lower skilled task of manual classification. This task is, however, not without its drawbacks, as human error can easily miss small areas, or misclassify water containing a lot of aquatic vegetation as land and vice versa. These latter ‘*missed*’ areas are of significance, as *Anopheles* mosquitoes are generally found in water containing vegetation. The fact that there was a 10% discrepancy in the manual classification undertaken by two independent researchers, both of whom were familiar with the study area, demonstrates the fallibility in this method.

As with the drone image capture, an alternative is to outsource these activities to an organisation which specialises in image processing and classification. Additionally, cloud-based computing services such as DroneDeploy’s Map Engine (DroneDeploy, 2018) and Google Earth Engine (Mutanga & Kumar, 2019) could be used as these allow individuals/groups to harness the power of remote servers to manage and manipulate the data. This approach could facilitate the development of more automated habitat classification approaches i.e. using data from other organisations, previous field or professional expertise in remote sensing to develop classification algorithms that don’t require the use of bespoke training data. The TropWet tool developed by Hardy, Oakes, & Ettritch (2020) is a demonstration of this in which satellite imagery (Landsat, 30m resolution) is automatically classified for a user-specified area and time period using a Google Earth Engine interface.

There are still practical challenges with these approaches, particularly relating to the upload of large image files to enable these processes to be undertaken remotely, however these may be preferable to the more technical challenge of managing the data in-house.

### Entomological survey

Larval surveys are an important part of the process of LSM both to confirm the species of mosquitoes found in the area, to characterise the types of surface water where larvae are likely to be found, and to monitor the progress of any subsequent intervention. Larval surveys are however a time-consuming process, particularly when undertaken during the wet season during which areas become inaccessible following heavy rains. The role of drones in LSM is not to completely remove the need for larval surveys, but to help differentiate between water bodies with respect to their potential as larval habitat and/or to differentiate sites according to their potential larval productivity.

We note that in our study that vegetation coverage appears to be important when considering presence/absence of late stage *Anopheles* larvae, with coverage correlated with the type of vegetation found i.e. coverage of floating vegetation was likely to be less than that of emergent or submerged vegetation. A full understanding the larval ecology of the local individual malaria vectors would greatly assist a targeted LSM approach aided by drone-imagery support. In south-eastern Tanzania, a basic characterisation of *An. funestus* larval habitats provides support that this species occupies small spring-fed pools, permanent natural ponds and slow-moving waters each of which fall under the ‘few, fixed and findable’ paradigm (Nambunga et al., 2020). In a recent study in Southern Malawi (Gowelo et al., 2020), *An. arabiensis* was the dominant species, with high densities being found in aquatic habitats surrounded by bare soil. A species-specific approach to identifying larval habitat using drone imagery may therefore be required, with imagery captured throughout the year to better understand the temporal dynamics of larval habitat and thereby optimise the impact of any potential intervention. These images could further be used to monitor the progress of LSM campaigns with, for example, a more accurate estimates of LSM coverage and demonstratable changes in the landscape because of habitat removal/modification. Further entomological surveillance remains pivotal to establish where and when LSM should be deployed and measure the impact of the intervention on malaria transmission potential.

## Conclusions

Our study demonstrates the potential for drone imagery to be used as a tool to support the identification of mosquito larval habitat in rural areas where malaria is endemic. While this technology has the capacity to complement the more labour-intensive approach of identifying larval habitat from the ground, there are technical challenges to overcome before it can be smoothly integrated into malaria control activities. We believe that outsourcing the capturing and processing of drone imagery to private companies with the equipment and skills necessary to extract the required information is a more practical approach to developing equivalent skills in house. These services are becoming increasingly available in other sectors such as agriculture, forestry and environmental monitoring and there are promising developments in the African drone sector to support this local capacity. We do however continue to emphasise that drone imagery should not be used to completely replace larval surveys. Instead we envisage that this technology could provide supplementary information which may help to reduce the time spent finding locations to be sampled, monitor environmental changes over time and help to guide the frequency and scale of any LSM intervention, ultimately increasing its potential for success. Further consultations between experts and stakeholders in the fields of drones, image analysis and vector control are needed to develop more detailed guidance on how this technology can be most effectively exploited.

## Author contributions

MCS and CMJ conceived the ideas and designed methodology; all authors collected the data; MCS, PK and KZ analysed the data; MCS and CMJ led the writing of the manuscript. All authors contributed critically to the drafts and gave final approval for publication.

## Data availability

Entomological data can be accessed at: https://figshare.com/s/25102b084f56f41a1ca8 All drone imagery referenced in this paper has been uploaded to OpenAerialMap from which it can be freely downloaded (https://map.openaerialmap.org/#/33.489675521850586,-13.050633215031182,13/user/5f1fe6f357ddda00054a0647?_k=mmgy4y).

