## Supplementary Information for "The application of drones for mosquito larval habitat identification in rural environments: a practical approach for malaria control?"

**Maladrone paper – Supplementary Information**

Survey locations

Table S1: Locations of reservoirs around which the drone image capture and entomological surveys were focused. The area covered represents the size of the area captured in the drone imagery.

| Name | Longitude | Latitude | Area ($km^2$) |
|---|---|---|---|
| Chitete | 33.48779 | -13.05738 | 1.824 |
| Champhantha | 33.45770 | -13.05999 | 1.044 |
| Lori | 33.38845 | -13.08206 | 0.439 |
| Chimphoyo | 33.38340 | -13.11194 | 1.660 |
| Malangano | 33.43645 | -13.06844 | 1.757 |
| Farm 1 | 33.54726 | -12.99336 | 0.378 |
| Farm 2 | 33.55744 | -13.00070 | 0.439 |
| Farm 3 | 33.57166 | -12.99638 | 0.368 |

Table S2: Number of training and testing segments that were manually created for each land class.

| **Macroclass** | **Class** | **Segments** |
|---|---|---|
| Water | Open water | 150 |
| | Floating aquatic vegetation | 150 |
| | Submerged aquatic vegetation | 150 |
| | Emergent aquatic vegetation | 150 |
| | TOTAL | 600 |
| Land | Trees/bushes | 250 |
| | Grass | 250 |
| | Bare soil | 100 |
| | TOTAL | 600 |
| Man-made features | Iron roof (120) | 120 |
| | Rusted iron roof (120) | 120 |
| | Thatched roof (120) | 120 |
| | Concrete yard/road (120) | 120 |
| | Dirt roads/paths (120) | 120 |
| | TOTAL | 600 |

Table S3: A description of the variables that were derived from the images captured by the drones

| **Sensor required** | **Variable name** | **Derivation** |
|---|---|---|
| **Standard RGB camera** | RGB | Values of Red, Green and Blue reflectance |
| | Elevation | Photogrammetric methods within Agisoft Metashape |
| | Slope | Derived from elevation |
| | Brightness | Red+Green+Blue |
| | Normalised difference Turbidity Index (NDTI) | (Red-Green)/(Red+Green) |

| | | |
|---|---|---|
| | Haralick texture variables | Energy, Entropy, Correlation, Inverse Difference Moment, Inertia, Cluster Shade, Cluster Prominence,<br>Haralick Correlation. Derived using the HaralickTextureExtraction tool within OTB |
| **NIR sensor** | NIR | Value of NIR reflectance |
| **Red and NIR** | Normalised Difference Vegetation Index (NDVI) | (NIR-Red)/(NIR+Red)* |
| | Soil Adjusted Vegetation Index (SAVI) | 0.5(NIR-Red)/(NIR+Red+0.5) |
| **Green and NIR** | Normalised Difference Water Index (NDWI) | (Green-NIR)/(Green+NIR) |

*A modified version of this equation was used on the image captured by the Sentera sensor as per the manufacturer's instructions

Classification maps

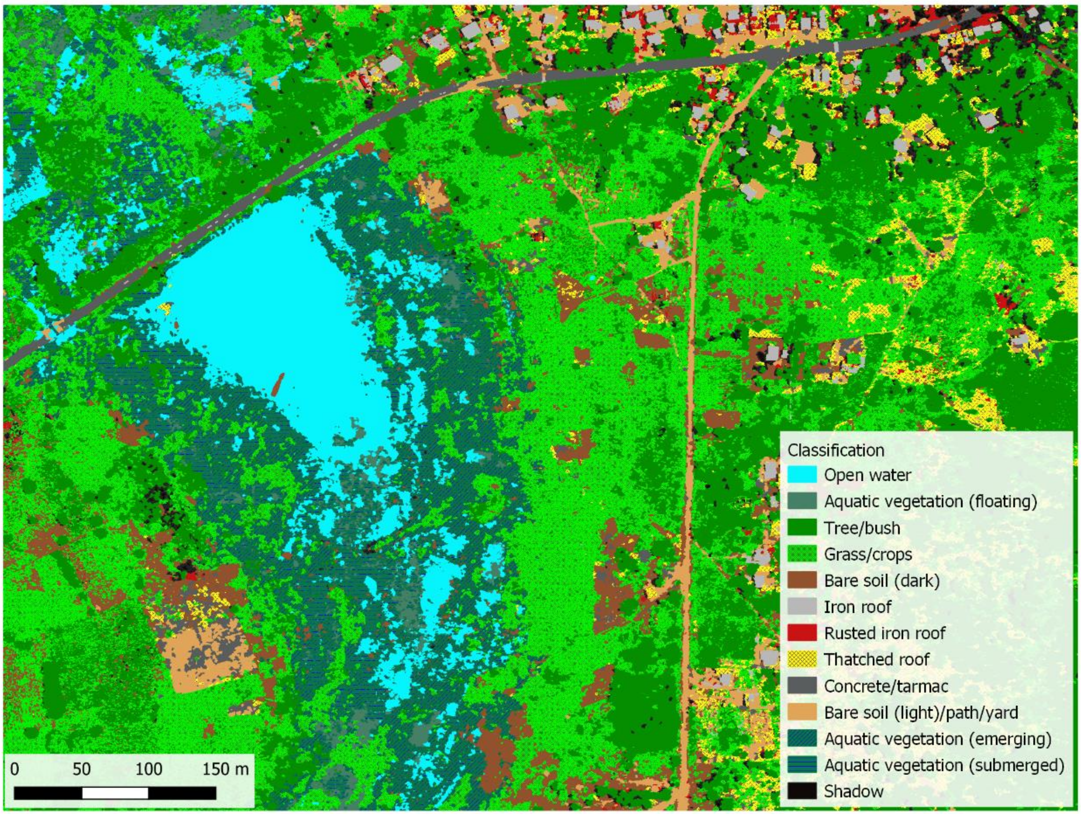

Figure S1: Example of a classification obtained using the random forests algorithm including NIR-derived variables.

Variable importance plots

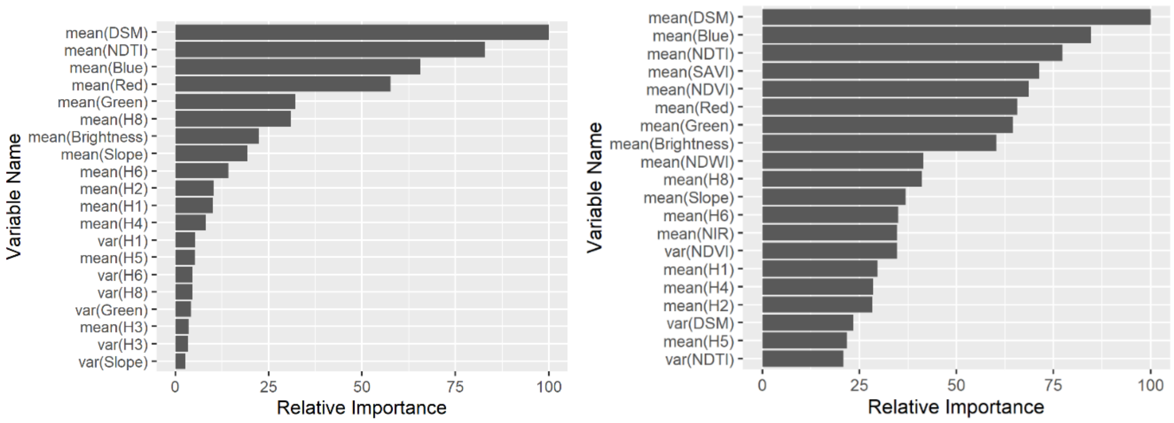

Figure S2: Variable importance plot corresponding to the classification presented in Figure 5 (without NIR) and Figure S1.

<u>Entomology data analysis</u>

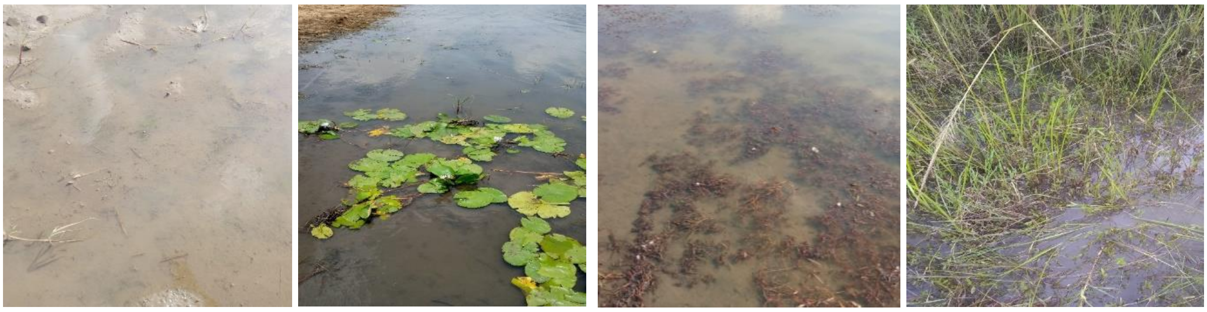

Figure S3: Examples of the vegetation types observed in the sampling sites – (from L to R) none, floating, submerged, emerging.

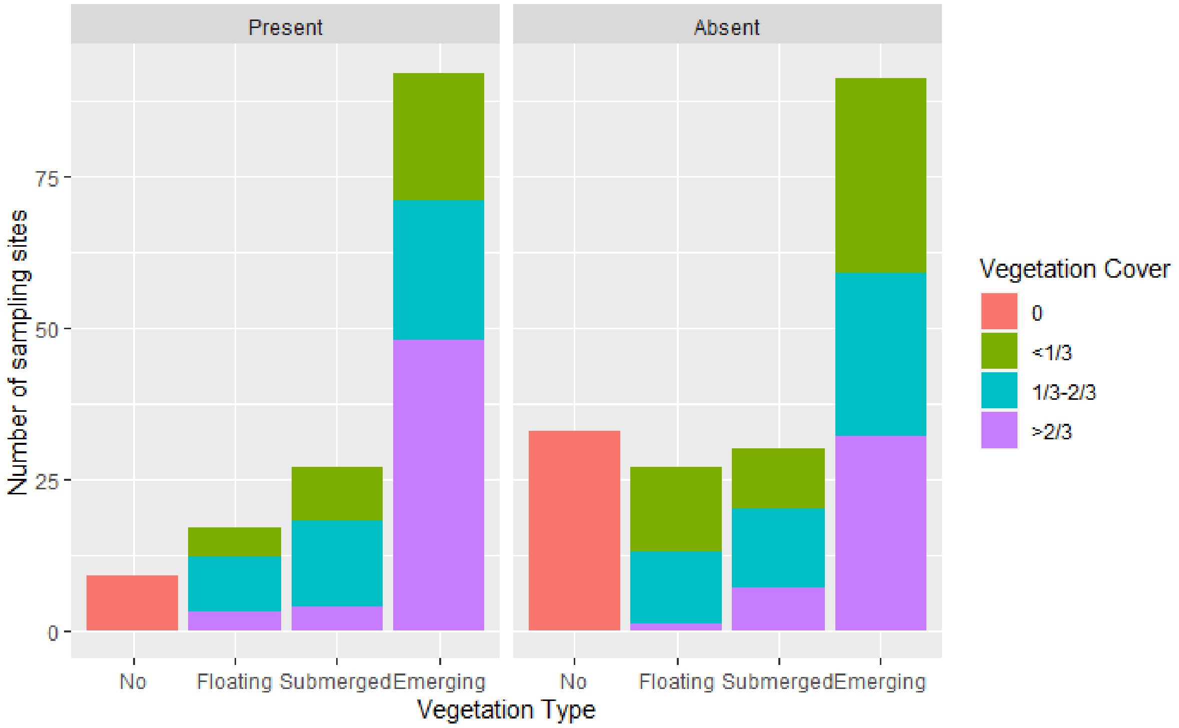

Figure S4: Summary of larval sampling sites by presence/absence of larvae, vegetation type and vegetation cover.

Table S4: Summaries of the larval sampling sites by presence/absence of late stage mosquito larvae found.

| | | **Any L3-L4 Larvae** | | | | |
|---|---|---|---|---|---|---|
| | | **Absent** | | **Present** | | **Total** |
| | | **N** | **(%)** | **N** | **(%)** | **N** |
| **Sampling period** | **2018 (early dry season)** | 60 | (46) | 68 | (53) | 128 |
| | **2019 (late dry season)** | 151 | (90) | 17 | (10) | 168 |
| | **2020 (wet season)** | 12 | (40) | 18 | (60) | 30 |
| **Vegetation** | **Yes** | 187 | (66) | 97 | (34) | 284 |
| | **No** | 36 | (86) | 6 | (14) | 42 |

| | | | | | | |
|---|---|---|---|---|---|---|
| **Dominant vegetation type** | **None** | 36 | (86) | 6 | (14) | 42 |
| | **Floating** | 32 | (73) | 12 | (27) | 44 |
| | **Submerged** | 36 | (63) | 21 | (37) | 57 |
| | **Emerging** | 119 | (65) | 64 | (35) | 183 |
| **Vegetation cover** | **0** | 36 | (86) | 6 | (14) | 42 |
| | **<$1/3$** | 68 | (75) | 23 | (25) | 91 |
| | **$1/3$ - $2/3$** | 63 | (64) | 35 | (36) | 98 |
| | **>$2/3$** | 56 | (59) | 39 | (41) | 105 |
| **Turbidity** | **Turbid** | 136 | (69) | 61 | (31) | 197 |
| | **Clear** | 87 | (67) | 42 | (33) | 129 |
| **Total** | | 223 | (68) | 103 | (32) | 326 |
